# A data-driven drug repositioning framework discovered a potential therapeutic agent targeting COVID-19

**DOI:** 10.1101/2020.03.11.986836

**Authors:** Yiyue Ge, Tingzhong Tian, Suling Huang, Fangping Wan, Jingxin Li, Shuya Li, Hui Yang, Lixiang Hong, Nian Wu, Enming Yuan, Lili Cheng, Yipin Lei, Hantao Shu, Xiaolong Feng, Ziyuan Jiang, Ying Chi, Xiling Guo, Lunbiao Cui, Liang Xiao, Zeng Li, Chunhao Yang, Zehong Miao, Haidong Tang, Ligong Chen, Hainian Zeng, Dan Zhao, Fengcai Zhu, Xiaokun Shen, Jianyang Zeng

## Abstract

The global spread of SARS-CoV-2 requires an urgent need to find effective therapeutics for the treatment of COVID-19. We developed a data-driven drug repositioning framework, which applies both machine learning and statistical analysis approaches to systematically integrate and mine large-scale knowledge graph, literature and transcriptome data to discover the potential drug candidates against SARS-CoV-2. The retrospective study using the past SARS-CoV and MERS-CoV data demonstrated that our machine learning based method can successfully predict effective drug candidates against a specific coronavirus. Our *in silico* screening followed by wet-lab validation indicated that a poly-ADP-ribose polymerase 1 (PARP1) inhibitor, CVL218, currently in Phase I clinical trial, may be repurposed to treat COVID-19. Our *in vitro* assays revealed that CVL218 can exhibit effective inhibitory activity against SARS-CoV-2 replication without obvious cytopathic effect. In addition, we showed that CVL218 is able to suppress the CpG-induced IL-6 production in peripheral blood mononuclear cells, suggesting that it may also have anti-inflammatory effect that is highly relevant to the prevention immunopathology induced by SARS-CoV-2 infection. Further pharmacokinetic and toxicokinetic evaluation in rats and monkeys showed a high concentration of CVL218 in lung and observed no apparent signs of toxicity, indicating the appealing potential of this drug for the treatment of the pneumonia caused by SARS-CoV-2 infection. Moreover, molecular docking simulation suggested that CVL218 may bind to the N-terminal domain of nucleocapsid (N) protein of SARS-CoV-2, providing a possible model to explain its antiviral action. We also proposed several possible mechanisms to explain the antiviral activities of PARP1 inhibitors against SARS-CoV-2, based on the data present in this study and previous evidences reported in the literature. In summary, the PARP1 inhibitor CVL218 discovered by our data-driven drug repositioning framework can serve as a potential therapeutic agent for the treatment of COVID-19.

## 1. Introduction

The outbreak of the pneumonia named COVID-19 caused by the novel coronavirus SARS-CoV-2 (2019-nCoV) has infected over 110,000 people worldwide by 8th March, 2020. Apart from China, other countries or regions including South Korea, Iran, and Europe have reported a rapid increase in the number of COVID-19 cases, implying that this novel coronavirus has posed a global health threat. Under the current circumstance of the absence of the specific vaccines and medicines against SARS-CoV-2, it is urgent to discover effective therapies especially drugs to treat the resulting COVID-19 disease and prevent the virus from further spreading. Considering that the development of a new drug generally takes years, probably the best therapeutic shortcut is to apply the drug repositioning strategy (i.e., finding the new uses of old drugs) [1, 2, 3] to identify the potential antiviral effects against SARS-CoV-2 of existing drugs that have been approved for clinical use or to enter clinical trials. Those existing drugs with potent antiviral efficacy can be directly applied to treat COVID-19 in a short time, as their safety has been verified in principle in clinical trials.

In this study, we applied a data-driven framework that combines both machine learning and statistical analysis methods to systematically integrate large-scale available coronavirus-related data and identify the drug candidates against SARS-CoV-2 from a set of over 6000 drug candidates (mainly including approved, investigational and experimental drugs). Our *in silico* screening process followed by experimental validation revealed that a poly-ADP-ribose polymerase 1 (PARP1) inhibitor, CVL218, currently in Phase I clinical trial, may serve as a potential drug candidate to treat COVID-19. Our *in vitro* assays demonstrated that CVL218 can exhibit effective inhibitory activity against SARS-CoV-2 replication in a dose-dependent manner and with no obvious cytopathic effect. In addition, we found that in human peripheral blood mononuclear cells (PBMCs), CVL218 is able to suppress the CpG-induced production of IL-6, which has been reported previously to be of high relevance to the viral pathogenesis of COVID-19, especially for those intensive care unit (ICU) patients infected by SARS-CoV-2. Further *in vivo* pharmacokinetic and toxicokinetic studies in rats and monkeys showed that CVL218 was highly distributed in the lung tissue and no apparent sign of toxicity was observed, which makes it an appealing potential drug candidate for the treatment of the novel pneumonia caused by SARS-CoV-2 infection. Moreover, our molecular docking study suggested that CVL218 may bind to the N-terminal domain of nucleocapsid (N) protein of SARS-CoV-2, providing a possible mode of its antiviral action against SARS-CoV-2. Based on the data present in this study and previous known evidences reported in the literature, we also discussed several putative mechanisms of the anti-SARS-CoV-2 effects for CVL218 or other PARP1 inhibitors to be involved in the treatment of COVID-19. Overall, our results indicated that the PARP1 inhibitor CVL218 identified by our drug repositioning pipeline may serve as an effective therapeutic agent against COVID-19.

## 2. Results

### 2.1. Overview of our drug repositioning framework

The overview of our data-driven drug repositioning framework is shown in Figure 1A. We first constructed a virus related knowledge graph consisting of drug-target interactions, protein-protein interactions and similarity networks from publically available databases (Methods). Three different types of nodes (i.e., drugs, human targets and virus targets) within the knowledge graph were connected through edges describing their interactions, associations or similarities to establish bridges of information aggregation and knowledge mining. We then applied a network-based knowledge mining algorithm to predict an initial list of drug candidates that can be potentially used to treat SARS-CoV-2 infection (Figure 1B and Methods). Next, we further narrowed down the list of drug candidates with the previously reported evidences of antiviral activities based on the text mining results from the large-scale literature texts, which were derived through a deep learning based relation extraction method named BERE [4] (Figure 1C and Methods), followed by a minimum of manual checking. After that, we used the connectivity map analysis approach [5] with the gene expression profiles of ten SARS-CoV-infected patients [6] to further refine the list of drug candidates against SARS-CoV-2 (Figure 1D, Table 1, Table S1 and Methods). The above screening process revealed that a poly-ADP-ribose polymerase 1 (PARP1) inhibitor PJ-34 could potentially have the antiviral activities against SARS-CoV-2.

**Figure 1.**
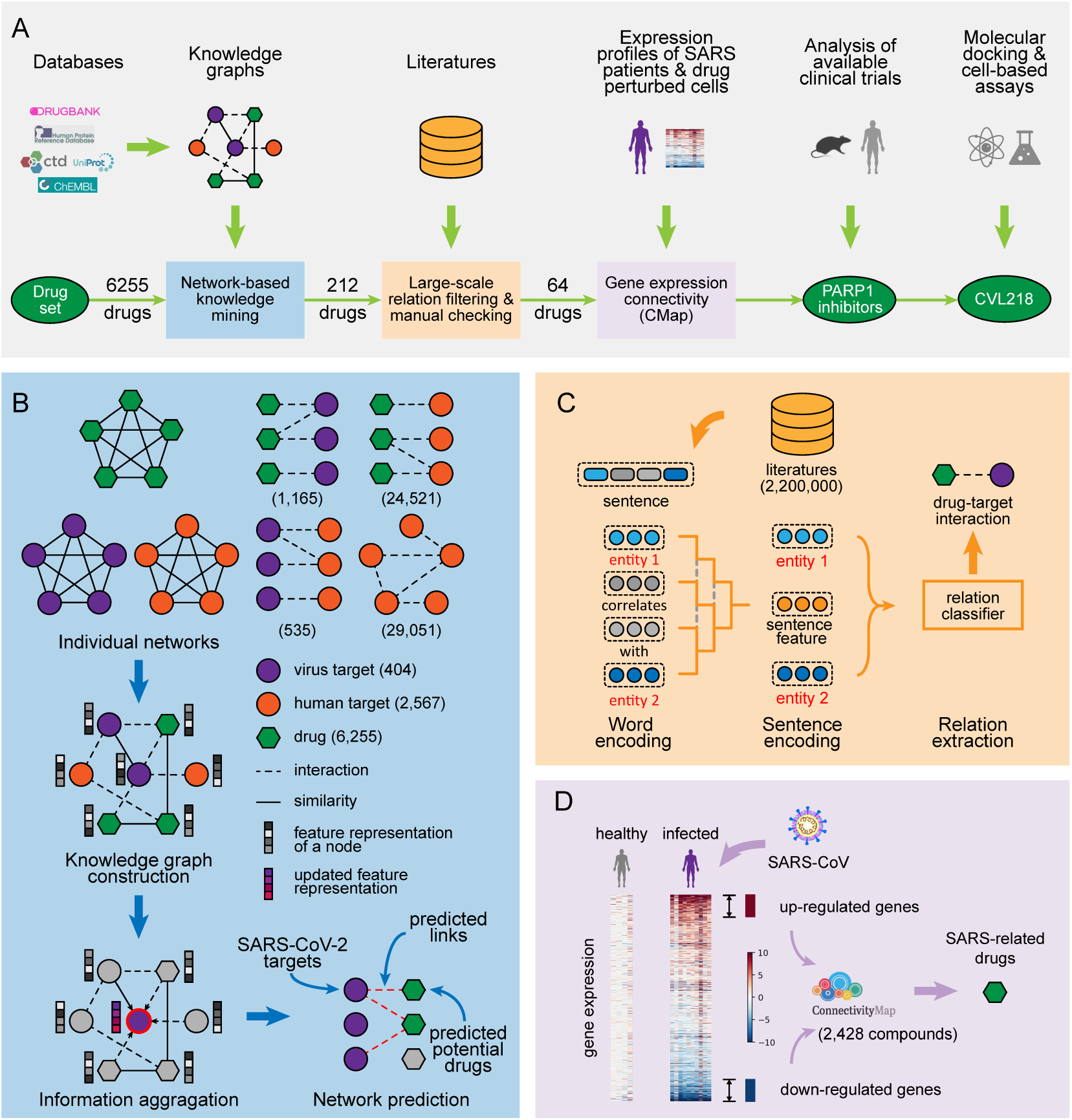
*(preceding page)*: Schematic illustration of our drug repositioning pipeline for discovering the potential drugs to treat the COVID-19 disease. (A). The overview of our drug screening pipeline. The initial drug set for screening contains 6255 drug candidates, mainly including 1786 approved drugs, 1125 investigational drugs and 3290 experimental drugs. The number of drug candidates after each filtering step is also shown. (B). The network-based knowledge mining module. Seven individual networks containing three types of nodes (i.e., drugs, human targets and virus targets) and the corresponding edges describing their interactions, associations or similarities are first constructed based on the known chemical structures, protein sequences and relations derived from publically available databases. Then a deep learning based method, which learns and updates the feature representation of each node through information aggregation, is used to predict the potential drug candidates against a specific coronavirus. (C). The automated relation extraction module. The structure of each sentence from the literature texts is first learned from the encoded word features using the Gumbel tree gated recurrent unit technique [4, 74]. Then the learned sequence structures as well as the corresponding encoded word features are fed into a relation classifier to automatically extract the relations between two entities from large-scale documents in the literature. (D). The connectivity map (CMap) analysis module. The transcriptome profiles of the Peripheral Blood Mononuclear Cell (PBMC) samples from the SARS-CoV infected patients and healthy persons are compared to derive the query gene expression signatures, which are then correlated to the drug-perturbed cellular expression profiles in the connectivity map [5] to filter out the anti-SARS-CoV drug candidates.

**Table 1:** The top list of drug candidates identified by the connectivity map analysis using the gene expression profiles of the peripheral blood mononuclear cell (PBMC) samples of ten SARS-CoV-infected patients [6]. The connectivity map score [5] of *−*90.0 was used as the cut-off threshold to determine the top list, i.e., only those drug candidates with the connectivity scores of the query ranked to the top 10% of the reference perturbations were selected. Two PARP1 inhibitors (i.e., veriparib and PJ-34) were chosen into the top list (shown in bold).

| Connectivity Map Score | Compound BRD ID | Name | Description |
| --- | --- | --- | --- |
| $-98.94$ | BRD-K87142802 | **Veliparib** | **PARP inhibitor** |
| $-95.37$ | BRD-A35338386 | NECA | Adenosine receptor agonist |
| $-95.26$ | BRD-K11853856 | **PJ-34** | **PARP inhibitor** |
| $-92.96$ | BRD-A53952395 | Prilocaine | Local anesthetic |
| $-91.8$ | BRD-K32977963 | Eugenol | Androgen receptor antagonist |
| $-91.56$ | BRD-A09495397 | Bicuculline | GABA receptor antagonist |
| $-91.32$ | BRD-K82164249 | Andarine | Androgen receptor modulator |

### 2.2. Validation of our network-based knowledge mining results

To demonstrate that our computational pipeline for drug repositioning can yield reasonably accurate prediction results, we also validated our network-based knowledge mining algorithm (Figure 1B) using the retrospective data of the two coronaviruses that are closely related to SARS-CoV-2 and had been relatively well studied in the literature, i.e., SARS-CoV and MERS-CoV. With the aid of our developed text mining tool BERE, we found that many of the drugs that had been reported previously in the literature to have antiviral activities against the corresponding coronavirus, were also among the top list of our predicted results (Table 2). For example, chloroquine, an FDA-approved drug for treating malaria [7], which was previously reported to exhibit micromolar anti-SARS-CoV activity *in vitro* [8], was also repurposed for targeting the same virus by our prediction framework. Gemcitabine, which was originally approved for treating certain types of cancers [9], was also predicted for targeting SARS-CoV with validation by previous *in vitro* studies [10]. Cyclosporine, a calcineurin inhibitor approved as an immunomodulatory drug [11], was observed to block the replication of SARS-CoV [12], and also successfully predicted by our approach. Among the predicted top list for MERS-CoV, miltefosine, which was approved for treating leishmaniasis [13], was previously identified to have anti-MERS-CoV activity [14]. Chlorpromazine and imatinib, which were used for treating schizophrenia [15] and leukemia [16], respectively, were also selected by our computational pipeline as anti-MERS-CoV drugs and can be validated by previous *in vitro* experiments [10]. Thus, the above retrospective study illustrated that our computational framework is able to predict effective drug candidates against a specific coronavirus.

**Table 2:** Selected examples of the predicted drug candidates against SARS-CoV or MERS-CoV that can be validated by the literature evidences in a retrospective study. The drug candidates were first predicted using our network based knowledge mining algorithm with a cut-off threshold of p-value *<* 0.05. Then the identified drug candidates were validated using an automated relation extraction method from the large-scale literature texts, followed by a minimum of manual checking.

| Drug name | Virus | Original targets<sup>a</sup> | Original indications<sup>b</sup> | References<sup>c</sup> |
| --- | --- | --- | --- | --- |
| Chloroquine | SARS-CoV | Fe(II)-protoporphyrin IX (Plasmodium falciparum) | Malaria | [8, 82] |
| Gemcitabine | SARS-CoV | Ribonucleoside-diphosphate reductase large subunit (Human) | Cancer | [10, 82] |
| Cyclosporine | SARS-CoV | Calcineurin subunit B type 2 (Human) | Prophylaxis of organ rejection, severe active rheumatoid arthritis (RA) | [12] |
| Indomethacin | SARS-CoV | Prostaglandin G/H synthase 1 and 2 (Human) | Symptomatic management of rheumatoid arthritis | [83] |
| Curcumin | SARS-CoV | Peroxisome proliferator-activated receptor gamma (Human) | Various pro-inflammatory diseases [84] | [85] |
| PJ-34 | SARS-CoV | Poly-ADP-ribose polymerase 1 (Human) | Experimental allergic encephalomyelitis [86] | [87] |
| Hesperetin | SARS-CoV | Sterol O-acyltransferase 1 (Human) | Lowering cholesterol [88] | [89] |
| Miltefosine | MERS-CoV | P-glycoprotein 1 (Human) | Mucosal, cutaneous, visceral leishmaniasis | [14] |
| Chlorpromazine | MERS-CoV | Dopamine D2 and D1 receptors, 5-hydroxytryptamine receptor 1A and 2A, Alpha-1A and -1B adrenergic receptors, Histamine H1 receptor (Human) | Schizophrenia and other psychotic disorders | [10, 82] |
| Imatinib | MERS-CoV | BCR-ABL fusion kinase (Human) | Leukemia | [82, 90] |

a. The parenthesis indicates the organism of the target(s).
b. Drug indications stand for the official indications approved by the FDA, obtained from DrugBank [17], unless other references are stated.
c. References stand for the supporting literatures.

### 2.3. CVL218 exhibits in vitro inhibitory activity against SARS-CoV-2 replication

As PJ-34 is still currently in the pre-clinical trial stage (DrugBank ID: DB08348, [17]), we selected two PARP1 inhibitors, including olaparib and mefuparib hydrochloride (CVL218) (Figure S1), that are currently FDA-approved and at Phase I clinical trial, respectively, for our initial study. We first conducted a pilot experimental test *in vitro* (Methods) on the anti-SARS-CoV-2 activities of olaparib, CVL218 and several other related drugs (Figure 2A). We found that both PARP1 inhibitors olaparib and CVL218 exhibited inhibitory effects against SARS-CoV-2 replication. Nevertheless, CVL218 showed a much higher inhibition rate than olaparib. More specifically, olaparib inhibited SARS-CoV-2 replication by 15.48% at a concentration of 3.2 *µ*M, while CVL218 reached 35.16% reduction at a concentration of 3 *µ*M.

**Figure 2.**
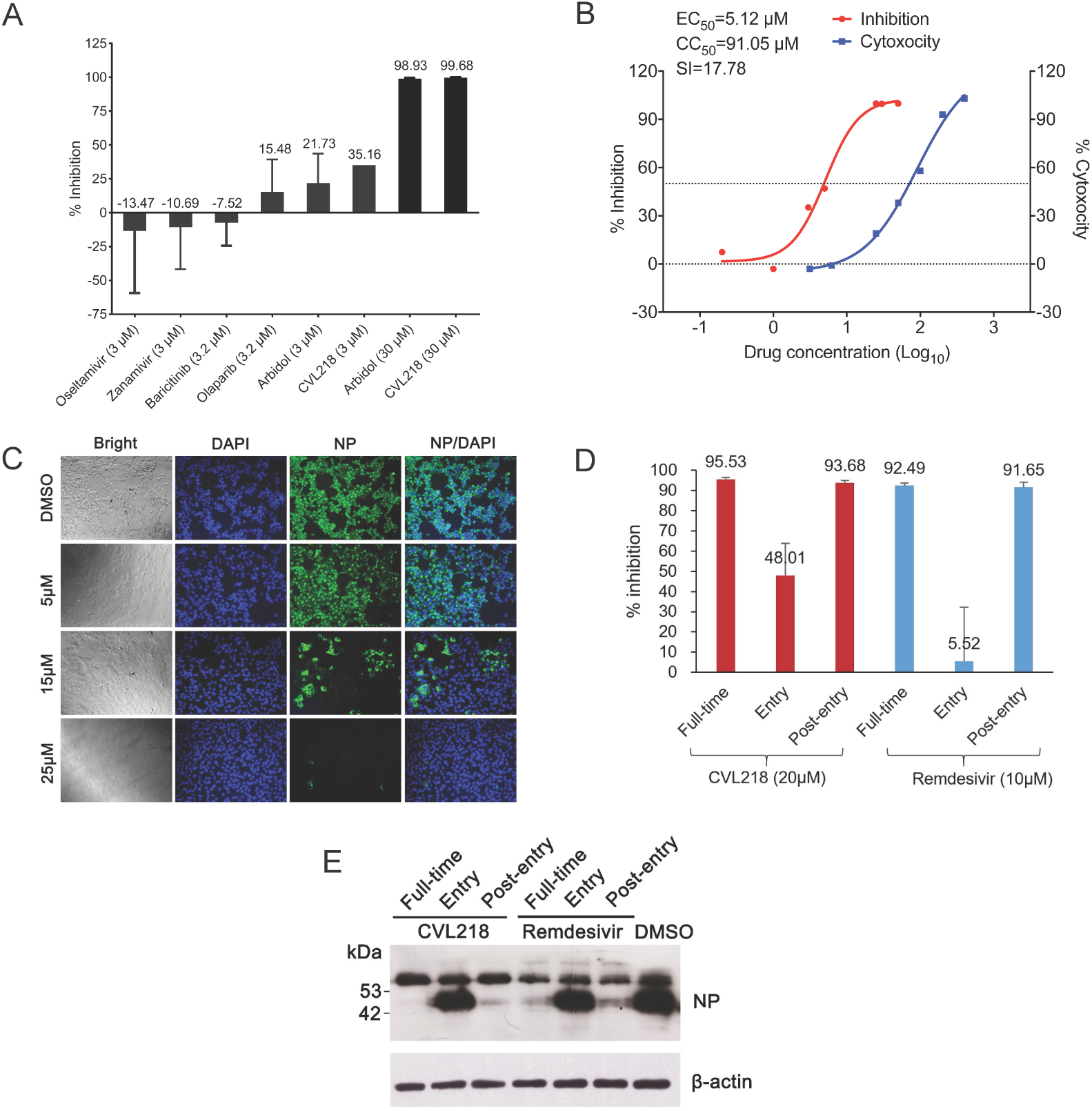
*(preceding page)*: The *in vitro* anti-SARS-CoV-2 activities of the tested drugs. (A). The *in vitro* inhibition rates of multiple tested drugs on SARS-CoV-2 replication at individual concentrations. (B). The concentration-dependent inhibition curve of CVL218 against SARS-CoV-2 replication and its cytotoxicity results. (C). Visualization of virus nucleoprotein (NP) expression of the infected cells upon treatment of CVL218 at 14 h post the SARS-CoV-2 infection using fluorescence microscopy. (D). Time-of-addition results on the inhibition of CVL218 and remdesivir against SARS-CoV-2 *in vitro*. The viral inhibitory activities of CVL218 and remdesivir were measured at “full-time”, “entry”, and “post-entry” stages, respectively. (E). Virus NP expression in the infected cells upon the treatment of CVL218 and remdesivir was analyzed by Western blot.

Notably, the antiviral efficacy of CVL218 even surpassed arbidol, which is one of the standard treatments for COVID-19 in the Diagnosis and Treatment Protocol for Novel Coronavirus Pneumonia (Trial Version 6) promulgated by the Chinese government (http://www.nhc.gov.cn/yzygj/s7653p/202002/8334a8326dd94d329df 351d7da8aefc2/files/b218cfeb1bc54639af227f922bf6b817.pdf). In particular, arbidol inhibited SARS-CoV-2 replication by 21.73% at 3 *µ*M, much lower than that of CVL218 at the same concentration (Figure 2A). In contrast, oseltamivir, zanamivir (drugs used for preventing influenza virus infection) and baricitinib (JAK1/2 inhibitor, which was recommended in [18] to treat COVID-19) showed no inhibitory activities against SARS-CoV-2 at the concentration of 3 *µ*M or 3.2 *µ*M.

Based on the above pilot experimental results, we then chose CVL218 for subsequent experimental studies. Our further *in vitro* assays (Methods) showed that CVL218 exhibited effective inhibitory activity against SARS-CoV-2 replication in a dose-dependent manner, with an EC_50_ of 5.12 *µ*M (Figure 2B). We also assessed the cytotoxicity of CVL218 by the CCK8 assay (Methods), and found that CVL218 had a CC_50_ of 91.05 *µ*M in Vero E6 cells. In addition, immunofluorescence microscopy (Methods) revealed that, at 14 h post SARS-CoV-2 infection, virus nucleoprotein (NP) expression in the CVL218-treated cells demonstrated a dose-response relationship with the treated drug concentrations, and was significantly lower upon CVL218 treatment compared with that in the DSMO treated cells (Figure 2C). No obvious cytopathic effect was observed in the infected cells treated with CVL218 at 25 *µ*M.

To systematically assess the inhibitory activities of CVL218 against SARS-CoV-2, we also performed a time-of-addition assay (Methods) to determine at which stage CVL218 inhibits viral infection. Remdesivir, which has entered the clinical trials for the treatment of COVID-19 (https://clinicaltrials.gov/ct2/show/NCT0425765 6), was also tested in this assay for comparison. In particular, as compared to the DMSO control group, both CVL218 and remdesivir showed potent antiviral activities during the full-time procedure of the SARS-CoV-2 infection in Vero E6 cells (Figure 2D). The results of the time-of-addition assay indicated that CVL218 can partially work against the viral entry and significantly inhibit the replication of virus post-entry, while the remdesivir can only function at the post-entry stage (Figure 2D, 2E). All together, the results of these *in vitro* assays indicated that CVL218 can be further evaluated as a potential therapeutic agent for treating COVID-19.

### 2.4. CVL218 inhibits IL-6 production in PBMCs induced by CpG-ODN 1826

Recently it has become evident that interleukin-6 (IL-6) is one of the most important cytokines during viral infection [19], and emerging clinical studies in humans and animals have linked the excessive synthesis of IL-6 with the persistence of many viruses, such as human immunodeficiency virus (HIV) [20], foot and mouth disease virus [21] and vesicular stomatitis virus (VSV) [22]. In addition, an *in vivo* study in the Friend retrovirus (FV) mouse model showed that IL-6 blockage can reduce viral loads and enhance virus-specific CD8+ T-cell immunity [23]. These findings supported a hypothesis that rapid production of IL-6 might be a possible mechanism leading to the deleterious clinical manifestations in viral pathogenesis [24]. Recently published researches on the clinical characteristics of severe patients with SARS-CoV-2 infection showed that IL-6 was significantly elevated especially in those ICU patients, which caused excessive activated immune response [25, 26, 27, 28, 29]. The pathological role of IL-6 in SARS-CoV-2 infection indicated that IL-6 blockade may provide a feasible therapy for the treatment of COVID-19.

To test whether CVL218 is able to regulate the IL-6 production *in vitro*, we stimulated the IL-6 production of the peripheral blood mononuclear cells (PMBCs) by CpG-ODN 1826, which is an effective stimulator of cytokines and chemokines. Incubation of PBMCs with 1 *µ*M CpG-ODN 1826 for 6 h (Methods) induced IL-6 production by 40%, when compared to untreated cells (Figure 3). In the presence of CVL218, the stimulatory effect of CpG-ODN 1826 was counteracted. Further study showed that CVL218 inhibited the CpG-induced IL-6 upregulation in a time- and dose-dependent manner (Figure 3). More specifically, exposure with CVL218 at concentrations 1 *µ*M and 3 *µ*M for 12 h attenuated the CpG-induced IL-6 production by 50% and 72.65%, respectively. These results provided an *in vitro* evidence to support CVL218 as a potential therapeutic agent for treating pro-inflammatory response caused by SARS-CoV-2 infection.

**Figure 3:**
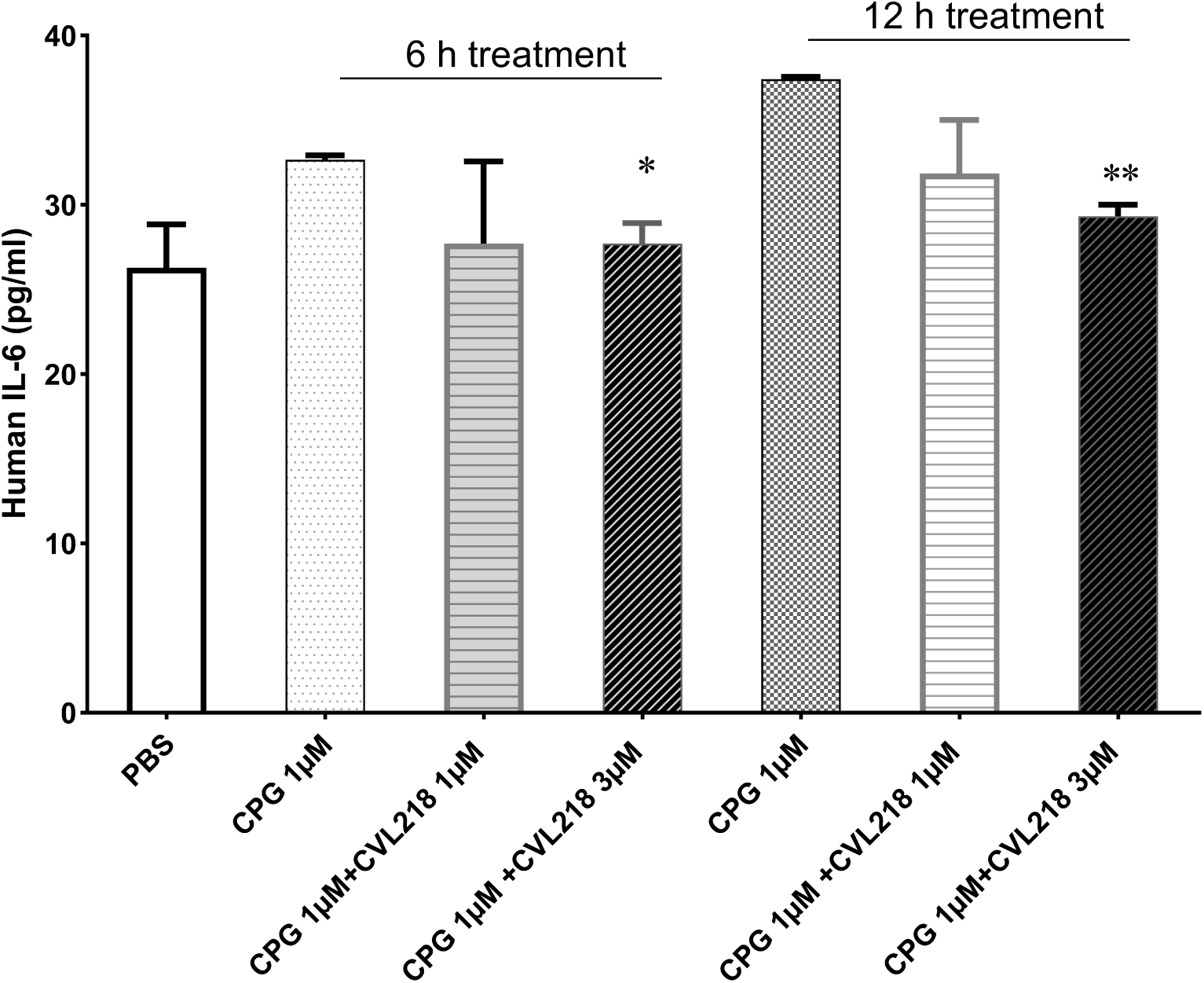
CVL218 attenuates the CpG-induced IL-6 production in a time- and dose-dependent manner.

### 2.5. CVL218 possesses good pharmacokinetic and toxicokinetic characteristics in animals

#### 2.5.1. CVL218 has the highest tissue distribution in lung of rats

We further performed *in vivo* pharmacokinetic and toxicokinetic evaluation of CVL218 in animals (Methods). We first examined the concentrations of CVL218 over different tissues in rats at different time points post oral administration at different doses (Figure S2 and Table S2), which was also previously reported in [30]. Among seven tissues (i.e., lung, spleen, liver, kidney, stomach, heart and brain), we observed that lung had the highest CVL218 concentration, which was 188-fold higher compared to that of plasma (Table 3). The observation that lung had the highest concentration of CVL218 was in line with the fact that the SARS-CoV-2 virus has the most pathological impact in lung with high viral loads, which suggested that CVL218 has the potential to be used for the indications of the lung lesions caused by SARS-CoV-2 infection, if its antiviral profile can be established in animal models and clinical trials.

**Table 3:** Comparison of the tissue to plasma concentration ratios between CVL218 and arbidol in rats. The concentrations of CVL218 over different tissues of rats were measured at the 180 min time point following 20 mg/kg oral administration. The concentrations of arbidol over different tissues of rats at the 15 min time point following 54 mg/kg oral administration were obtained from the literature [91, 92]. Means and standard deviations are shown.

| Tissue | Tissue to plasma concentration ratio |  |  |  |
| --- | --- | --- | --- | --- |
|  | CVL218 |  | Arbidol |  |
| Lung | 188.364 | ± 28.467 | 0.553 | ± 0.392 |
| Spleen | 54.897 | ± 6.250 | 0.110 | ± 0.060 |
| Liver | 46.780 | ± 5.215 | 0.204 | ± 0.062 |
| Kidney | 41.307 | ± 5.391 | 0.055 | ± 0.040 |
| Stomach | 22.133 | ± 7.130 | 4.920 | ± 2.159 |
| Heart | 16.514 | ± 1.348 | 0.028 | ± 0.015 |
| Brain | 9.728 | ± 1.130 | 0.011 | ± 0.002 |

Furthermore, we compared the pharmacokinetic data between CVL218 and arbidol, a broad-spectrum antiviral drug commercialized in China since 2016, had been recommended to treat the SARS-CoV-2-infected patients by the Chinese government. We found that the pharmacokinetic parameters of CVL218 and arbidol were comparable, with similar plasma concentrations and drug exposures (Table S3). Arbidol was mostly distributed in stomach and plasma post administration in rats. In contrast, higher distributions of CVL218 in tissues especially in lung rather than plasma compared to those of arbidol indicated a superior pharmacokinetic profile of CVL218, which may render it as a better potential antiviral treatment of SARS-CoV-2 infection in lung.

#### 2.5.2. The toxicity study demonstrated a safety profile of CVL218 in rats

In rats after being orally administrated 20/60/160 mg/kg of CVL218 for 28 consecutive days and followed by 28 more days without drug administration (Methods), we observed no significant difference in body weight of rats among different dosage and the control groups (Figure 4A).

**Figure 4:**
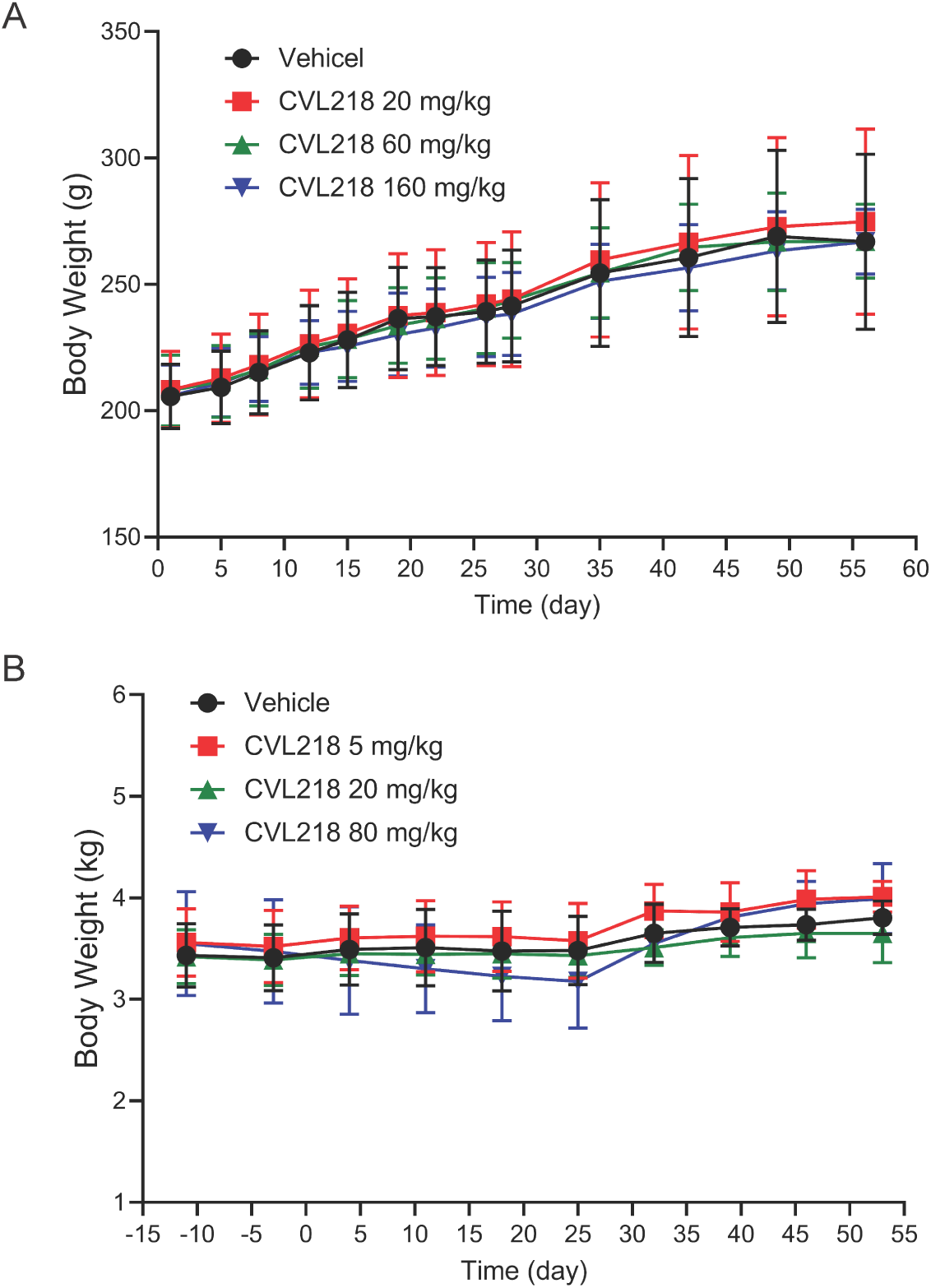
Effects of CVL218 on body weight in rats (A) and monkeys (B). Rats and monkeys were orally administered 20/60/160 mg/kg and 5/20/80 mg/kg of CVL218, respectively, for 28 consecutive days and then followed by 28 more days without drug administration.

We next conducted a toxicokinetic analysis of CVL218 in rats (Methods). In particular, rats were given CVL218 20/60/160 mg/kg by oral gavage once a day for consecutive 28 days, followed by 28 days without CVL218 administration, to investigate the reversibility of the toxic effects of the compound and examine whether there is any potential delayed-onset toxicity of this drug in rats. The results showed that, the maximum tolerable dose (MTD) and the no-observed adverse effect level (NOAEL) were 160 mg/kg and 20 mg/kg, respectively. The exposure of female rats to CVL218 (AUC_0_*_−_*_24_) was 7605 h*·*ng/mL in day 1 and 6657 h*·*ng/mL in day 28, while that of male rats (AUC_0_*_−_*_24_) was 9102 h*·*ng/mL in day 1 and 10253 h*·*ng/mL in day 28 (Table S4). Based on the toxicokinetic results from the repeated dose studies, all rats survived after a 28-day treatment period and showed no apparent signs of toxicity.

#### 2.5.3. CVL218 exhibits a favorable safety profile in monkeys

Monkeys were administered CVL218 (5, 20 or 80 mg/kg) by nasogastric feeding tubes with a consecutive daily dosing schedule for 28 days, followed by a 28-day recovery period (Methods). Only a slight decrease of body weight was observed in the high-dose (80 mg/kg) group, and all changes were reversed after a 28-day recovery period (Figure 4B), demonstrating a favorable safety profile for CVL218 in monkeys. Further examination of the toxicokinetic data of CVL218 in monkeys showed that the increase of the exposure of CVL218 (AUC_0_*_−_*_24_) was approximately dose proportional, and after consecutive 28 days of drug administration, the accumulation was not apparent. The exposure of female monkeys to CVL218 (AUC_0_*_−_*_24_) was 19466 h*·*ng/ml in day 1 and 18774 h*·*ng/ml in day 28 (Table S5), while that of male monkeys (AUC_0_*_−_*_24_) was 16924 h*·*ng/ml in day 1 and 22912 h*·*ng/ml in day 28. The maximum tolerable dose (MTD) of CVL218 in monkeys was 80 mg/kg, and the dose of 5 mg/kg was considered as the no-observed adverse effect level (NOAEL).

Overall, the above *in vivo* data showed that CVL218 possesses good pharmacokinetic and toxicokinetic characteristics in rats and monkeys, and its high-level distribution in the therapeutically targeted tissue (i.e., lung) may greatly favor the treatment of SARS-CoV-2 infection.

### 2.6. Molecular docking suggests the interactions between PARP1 inhibitors and the N-terminal domain of coronavirus nucleocapsid protein

As the previous studies have reported that the PARP1 inhibitor PJ-34 can target the N-terminal domain (NTD) of the coronavirus nucleocapsid (N) protein to reduce its RNA binding and thus impede viral replication [31, 32, 33], we speculated that olaparib and CVL218 may also interact with the N protein of SARS-CoV-2 to perform the similar antiviral function. To test this hypothesis, we conducted molecular docking to study the potential interactions between these two drugs and the N-NTD of SARS-CoV-2 (Methods).

The overall structure of HCoV-OC43 (another coronavirus phylogenetically closely related to SARS-CoV-2) and SARS-CoV-2 share similar compositions of secondary structure elements, including five conserved *β* strands and flexible loops (Figure 5A). In addition, their sequences covering the corresponding binding pocket regions are well conserved (Figure 5C). Thus, their structures may provide common molecular features in terms of interactions with small molecules at this binding pocket. Therefore, we also used the experimentally solved structure of HCoV-OC43-N-NTD complexed with PJ-34 as a reference to analyze our docking results.

**Figure 5.**
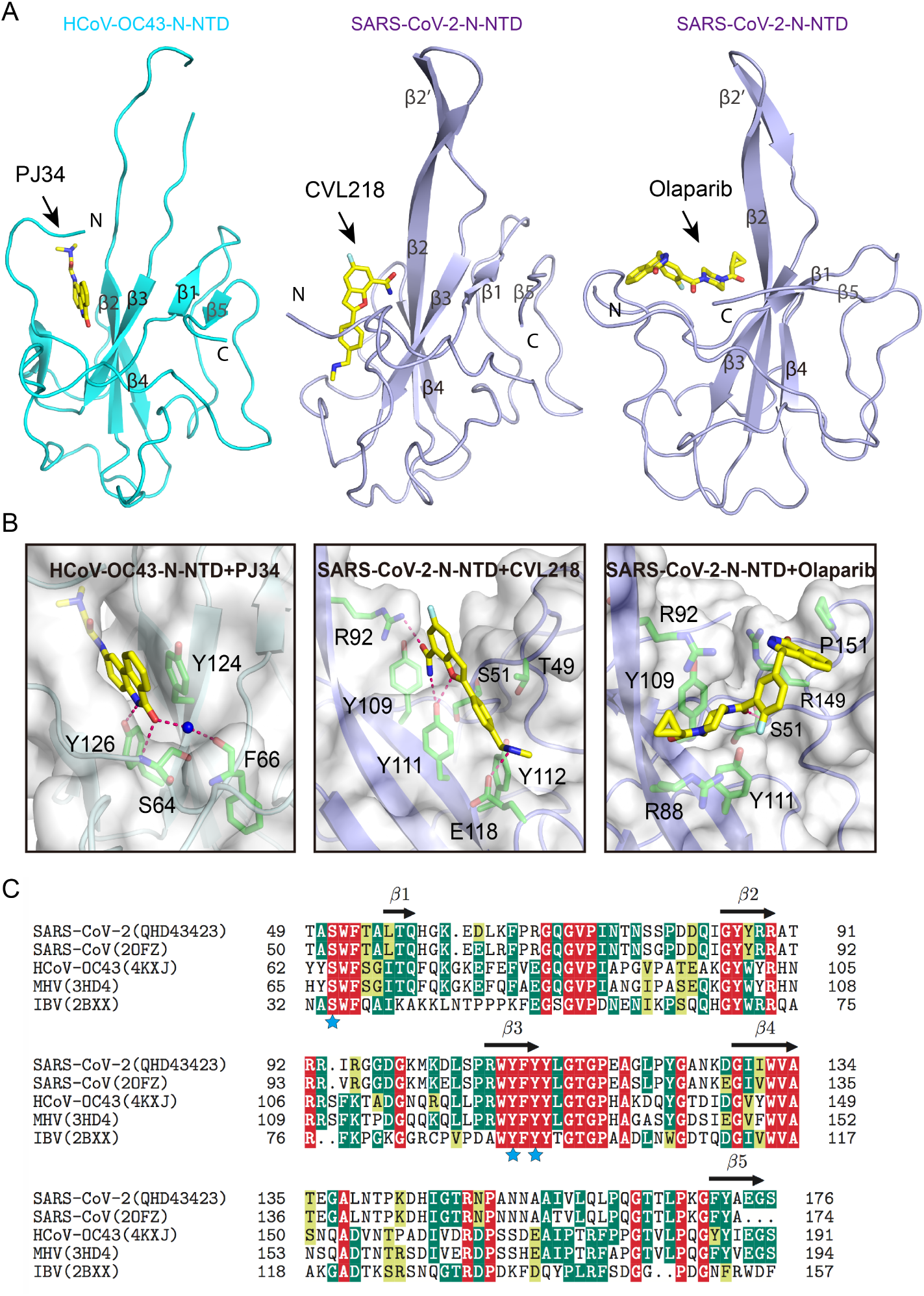
*(preceding page)*: The modeled structure of the N-terminal domain of nucleocapsid protein (N-NTD) of SARS-CoV-2 complexed with PARP1 inhibitors. (A). Overall complex structures of HCoV-OC43-N-NTD (i.e., the N-NTD of HCoV-OC43) bound to PJ-34 (PDB ID: 4kxj), and SARS-CoV-2-N-NTD (i.e., the N-NTD of SARS-CoV-2) bound to CVL218 and olaparib (both modeled by AutoDock4.2) in ribbon view. The structure of SARS-CoV-2-N-NTD that we used for docking simulation was derived from homology modeling [80], with residues ranging from 47 to 177 (GenBank: QHD43423). (B). Detailed molecular interactions between the coronavirus N-NTDs and PARP1 inhibitors. Left panel: The experimentally solved complex structure of HCoV-OC43-N-NTD (cyan ribbon) bound to PJ-34 (yellow sticks). Middle panel: The modeled complex structure of SARS-CoV-2-N-NTD (purple ribbon) bound to CVL218 (yellow sticks). Right panel: The modeled complex structure of SARS-CoV-2-N-NTD (purple ribbon) bound to olaparib (yellow sticks). The key residues interacting with the inhibitors are shown as green sticks. The hydrogen bonds are denoted as pink dashes. (C). Multiple sequence alignment (performed using MUSCLE [81]) of the N-NTDs among SARS-CoV-2, SARS-CoV, HCoV-OC43, mouse hepatitis virus (MHV) and infectious bronchitis virus (IBV). The virus names are listed on the left with available PDB codes shown in the parentheses. The sequence of SARS-CoV-2-N-NTD was obtained from GenBank (QHD43423). Secondary structure elements of HCoV-OC43-N are depicted above the sequence alignment. Asterisks indicate the key residues interacting with the inhibitors. The residues conserved among all five viruses are shaded in red, the residues with the percentage of conservation larger than 50% are shaded in green, and the similar residues are shaded in yellow.

Our docking results (more details can also be found in Supplementary Materials) showed that both CVL218 and olaparib can bind to the N-NTD of SARS-CoV-2 around the same binding pocket as in the experimentally solved complex structure between PJ-34 and the corresponding protein of HCoV-OC43, though with different binding poses (Figure 5B). Examination of docked structures indicated that CVL218 exhibits stronger binding ability than olaparib in terms of the hydrogen bond formation. Meanwhile, the key residues (i.e., S51, Y109 and Y111) participating in the binding with the drugs on SARS-CoV-2-N-NTD are also highly conserved among other viruses including SARS-CoV, HCoV-OC43, mouse hepatitis virus (MHV) and infectious bronchitis virus (IBV) (Figure 5C), suggesting that the N-NTDs of different viruses most likely display similar binding behaviors for PJ-34, CVL218 or other PARP1 inhibitors. Overall, our docking results indicated that CVL218 should be more effective in binding toward the nucleocapsid protein of SARS-CoV-2 compared to olaparib, thus better beneficial to intervene the nucleocapsid-dependent assembly of viral genome and thus inhibit viral replication.

## 3. Discussion

In this study we reported a top down data integration approach by combining both machine learning and statistical analysis techniques, followed by web-lab experimental validation, to identify potential drug candidates for treating SARS-CoV-2 infection. We showed that the PARP1 inhibitor CVL218 discovered by our *in silico* drug repurposing framework may have the therapeutic potential for the treatment of COVID-19. Although we mainly conducted *in vitro* assays to experimentally validate the anti-SARS-CoV-2 effects of olaparib and CVL218 due to limited time, it is natural to speculate that other PARP1 inhibitors may also have antiviral activities against SARS-CoV-2 infection, based on our computational prediction and experimental validation results.

Based on the data present in this study and the previously known evidences reported in the literature, we propose several potential mechanisms to help understand the involvement of PARP1 inhibitors in the treatment of COVID-19 (Figure 6). First, during the life cycle of the coronavirus, PARP1 inhibitors may inhibit the viral growth through suppressing viral replication and impeding the binding of the nucleocapsid protein to viral RNAs [31, 34, 35, 36], which can also be supported by our molecular docking results (see Section 2.6). Second, PARP1 inhibitors have been previously reported to play a critical role in regulating inflammatory response by modulating the expression of pro-inflammatory factors such as NF-*κ*B, AP-1, IL-6 and downstream cytokines and chemokines [37, 38, 39, 40]. Also, it has been shown that the overactivation of PARP1 promotes energy (NAD^+^/ATP) consumption and drives cell death, exacerbating the inflammation response [37, 38, 39, 41]. PARP1 inhibitors thus may repress the NF-*κ*B-mediated pro-inflammatory signals, and reduce energy failure and subsequent cell death induced by necrosis, thus providing a clinical potential for attenuating the cytokine storm and inflammatory response caused by SARS-CoV-2 infection. Third, ADP-ribosylation is a conserved post-translational modification on the nucleocapsid proteins across different coronavirus lineages, implying that it may have an important regulatory role for the structure packing of viral genome. Several previous studies have demonstrated that PARP1 is critical for viral replication [35, 42, 43]. For example, PARP1 has been reported to interact with hemagglutinin (HA) of influenza A virus (IAV) and promote its replication by triggering the degradation of host type I IFN receptor [44]. In addition, the ADP-ribosylation of adenoviral core proteins displays an antiviral defense mechanism [34]. Therefore, intervening the ADP-ribosylation mediated interplay between PARP1 and viral proteins may be another important pathway for PARP1 inhibitors to prevent SARS-CoV-2 infection. Of course, to throughly understand the anti-SARS-CoV-2 roles of PARP1 inhibitors, more experimental studies and direct (clinical) evidences will be needed in the future.

**Figure 6.**
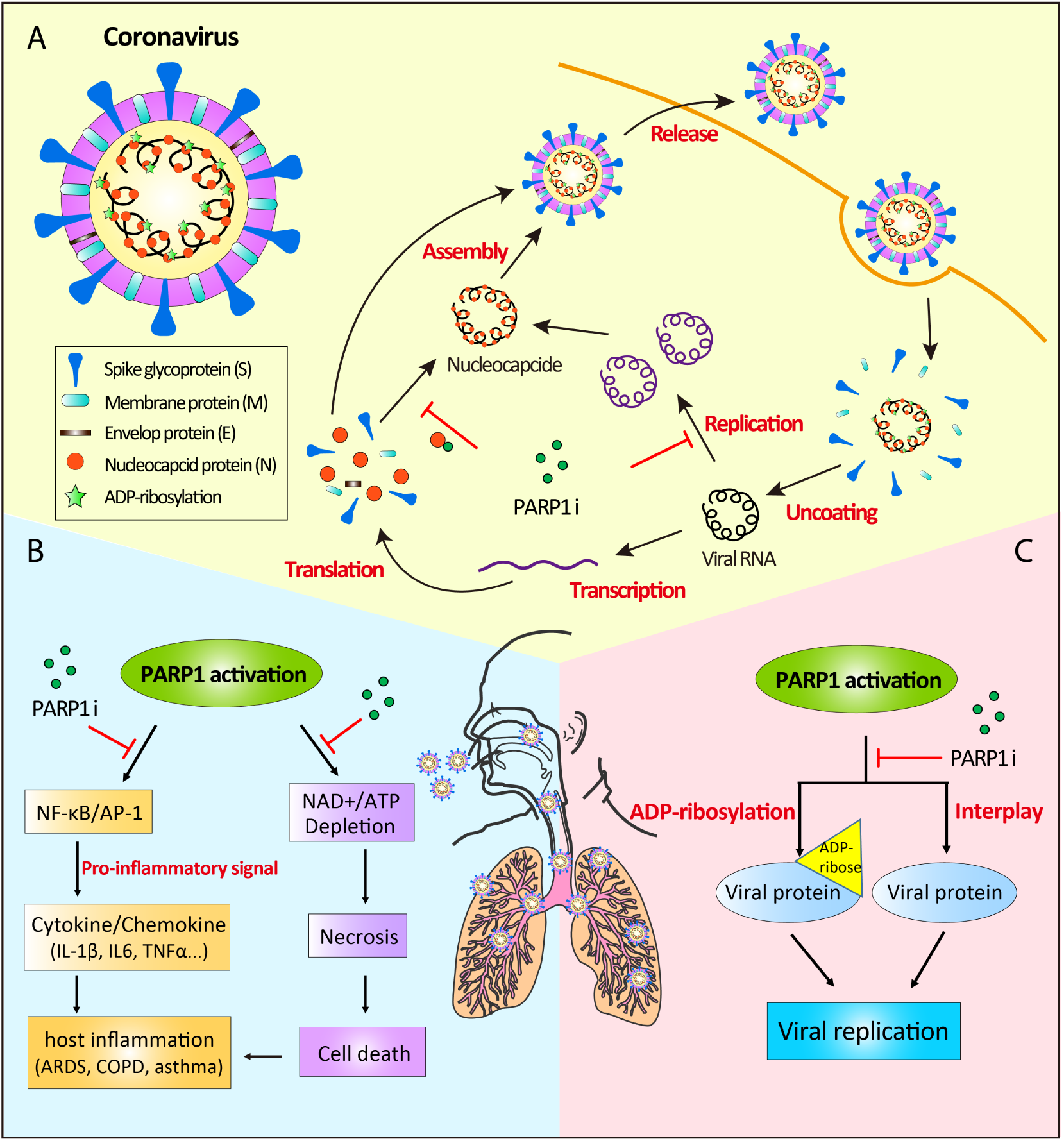
*(preceding page)*: The putative mechanisms for PARP1 inhibitors to combat the COVID-19 disease, derived based on the data present in this study and the known antiviral activities of PARP1 inhibitors previously reported in the literature. (A). Schematic diagram showing the possible antiviral mechanisms of PARP1 inhibitors in the life cycle of coronavirus in human cells. PARP1 inhibitors have been previously reported in the literature to suppress viral replication and imped the binding of nucleocapsid protein to viral RNAs, thus preventing the virus infection [31, 34, 35, 36]. (B). Potential protective effects of PARP1 inhibitors in the treatment of COVID-19. The antiinflammation effects of PARP1 inhibitors may be achieved through two possible molecular pathways. The first one is to modulate the expression of pro-inflammation factors such as NF-*κ*B, AP-1, IL-6 and downstream cytokines and chemokines [37, 38, 39, 40]. The second possible pathway is to prevent the overactivation of PARP1 and thus avoid the depletion of NAD^+^ and ATP, and the consequent cellular energy failure and cell death caused by necrosis [37, 38, 39, 40]. (C). The potential antiviral effects of PARP1 inhibitors through suppressing the ADP-ribosylation of viral proteins and intervening the host-pathogen interactions, thus resulting in the inhibition of viral replication [34, 35, 42, 43].

Considering the pro-inflammatory role of PARP1, the therapeutic effects of PARP1 inhibitors in inflammatory-mediated diseases have been extensively studied over past two decades [45, 46]. PJ-34, the early generation PARP1 inhibitor, has been suggested in previous studies to have neuroprotective effects in stroke model and protect mice from necroptosis-associated liver injuries by repressing the IL-33 expression [47, 48]. In addition, the FDA-approved PARP1 inhibitor, olaparib, has been reported to protect against the LPS (Lipopolysaccharide)-induced acute lung and kidney injuries in a NF-*κ*B-dependent manner in mice [49]. Numerous pre-clinical studies demonstrated that PARP1 inhibitors play an essential role in a range of inflammatory injuries and related diseases, especially the lung inflammatory disorders including ARDS (Acute Respiratory Distress Syndrome), COPD (Chronic Obstructive Pulmonary Disease) and asthma [40, 46, 50, 51]. All these studies suggest that PARP1 inhibitors are of high relevance to the treatment of the novel pneumonia caused by SARS-CoV-2 infection, possibly via their roles in modulating inflammatory response.

Notably, current pathological studies have shown that the severe patients infected by SARS-CoV-2 generally have higher plasma levels of IL-2, IL-6, IL-10, TNF*α*, IFN-*γ* [25, 27, 28, 29], implying a high risk of the inflammatory-associated cytokine storm after viral infection. In addition, reduction and functional exhaustion of T cells have also been observed in COVID-19 patients [27]. Therefore, blocking the overactive inflammatory response may be an effective strategy for the treatment of COVID-19, particularly for those ICU patients infected by SARS-CoV-2. Recently, tocilizumab, a monoclonal antibody drug targeting IL-6, has been recommended for the treatment of COVID-19 in the Diagnosis and Treatment Protocol for Novel Coronavirus Pneumonia (Trial Version 7) promulgated by the Chinese government (http://www.nhc.gov.cn/yzygj/s7653p/202003/46c9294a7dfe4cef80dc7f5912eb1989/files/ce3e6945832a438eaae415350a8ce964.pdf), which also highlights the vital role of anti-inflammatory response in current therapeutics against SARS-CoV-2. Our *in vitro* study has showed that CVL218 can effectively inhibit the IL-6 production induced by CpG in PBMCs (Figure 3). This finding indicates that CVL218 may also possess the IL-6 specific anti-inflammatory effect that is applicable to those severe patients infected by SARS-CoV-2.

PARP1 inhibitors are originally used for targeting homologous recombination repair defects in cancers, and mainly categorized as oncology drugs. Thus, it would generally need more safety data to justify any repurposing of PARP1 inhibitors for non-oncology indications. Fortunately, there are numerous existing pre-clinical and clinical studies on repurposing PARP1 inhibitors into non-oncological diseases, including the aforementioned acute diseases (e.g., acute respiratory distress syndrome (ARDS), stroke) [52] and chronic diseases (e.g., rheumatoid arthritis and vascular diseases) [52, 53]. All these evidences indicate the possibility of repurposing PARP1 inhibitors as a safe therapeutic agent to treat the current acute lung disease caused by SARS-CoV-2 infection. In addition, our pharmacokinetic and toxicokinetic data in rats and monkeys shown in our study indicate that CVL218 may have a relatively acceptable safety profile to be repositioned for the antiviral purpose. Moreover, CVL218 has been approved to enter Phase I clinical trial in 2017 by National Medical Products Administration (NMPA) in China for cancer treatment. The preliminary data from the Phase I clinical trial have shown that CVL218 is well tolerated in ascending dose studies at doses as high as 1000 mg QD and 500 mg BID, and no Grade II and above adverse events have been observed, which indicates that CVL218 is also quite safe and well tolerated in human. Our pharmacokinetic examination in rats has shown that CVL218 has the highest level distribution in the lung tissue, 188-fold higher concentration compared to that in plasma. Such a tissue specific enrichment in lung may bring an extra advantage for CVL218 to be used for the anti-SARS-CoV-2 purpose, as lung is the therapeutically targeted tissue for COVID-19. Moreover, high level distribution in lung may also suggest that only low dosage is needed in order to ensure the therapeutic efficacy of CVL218 against SARS-CoV-2, which may further reduce the risk of adverse events. Thus, CVL218 may have great potential to be repurposed as an effective therapeutic agent to combat SARS-CoV-2 and prevent future epidemic outbreak.

## 4. Methods

### 4.1. Construction of the virus-related knowledge graph

The virus-related knowledge graph was constructed for predicting the coronavirus related drugs. In total seven networks were considered in the constructed knowledge graph (Figure 1B), including a human target-drug interaction network, a virus target-drug interaction network, a human protein-protein interaction network, a virus protein-human protein interaction network, a drug molecular similarity network, a human protein sequence similarity network, and a virus protein sequence similarity network. The human target-drug interaction network was derived from DrugBank (version 5.1.0) [17]. The virus target-drug interaction network was constructed from the integrated data from DrugBank (version 5.1.0) [17], ChEMBL (release 26) [54], TTD (last update 11 Nov, 2019) [55], IUPHAR BPS (release 13, Nov, 2019) [56], BindindDB [57] and GHDDI (https://ghddi-ailab.github.io/Targeting2019-nCoV/CoV_Experiment_Data/), with a cut-off threshold of IC_50_/EC_50_/K*_i_*/K*_d_ <*10 *µ*M. The human protein-protein interaction network and the virus protein-human protein interaction network were constructed from the integrated data from BioGRID (release 3.5.181) [58], HuRI [59], Instruct [60], MINT (2012 update) [61], PINA (V2.0) [62], SignaLink (V2.0) [63] and innatedb [64]. The drug molecular similarity network was obtained by calculating the Tanimoto similarities from Morgan fingerprints with a radius of 2 computed using the rdkit tool [65]. The protein sequence similarity networks of both human and virus were obtained by calculating the Smith-Waterman similarities of the amino acid sequences derived from UniProt [66] using a sequence alignment software provided in [67]. Noted that we collected additional protein sequences of SARS-CoV-2 from UniProt [66] and added them into the corresponding networks for the final prediction. Those drugs without drug-target interactions or outside the Drug-Bank database were removed from the corresponding networks. We then constructed the virus-related knowledge graph by merging together all the nodes and edges of the above seven networks (Figure 1B). The constructed knowledge graph *G* = (*V, E*) is an undirected graph, in which each node *v ∈ V* in the node set *V* belongs to one of the node types (including drugs, human proteins, and virus proteins), and each edge *e ∈ E* in the edge set *E ⊂ V × V × R* belongs to one of the relation types from the relation type set *R* (including two drug-target interactions, two protein-protein interactions and three similarities).

### 4.2. The network-based knowledge mining algorithm

The initial list of drug candidates targeting SARS-CoV-2 was first screened using a network-based knowledge mining algorithm modified from our previous work [68, 69]. The goal was to capture the hidden virus-related feature information and accurately predict the potential drug candidates from the constructed knowledge graph, which was realized through learning a network topology-preserving embedding for each node.

More specifically, our model used a graph convolution algorithm [70] to gather and update feature information for each node in the constructed heterogeneous knowledge graph network from neighborhoods so that the network topology information can be fully exploited. Suppose that we perform *T* iterations of graph convolution. At iteration 1 *≤ t ≤ T*, the message 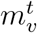 passed to node *v* can be expressed as:

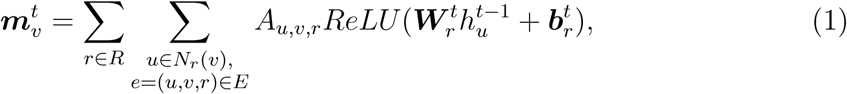

where *a_v,u,r_* stands for the weight for edge 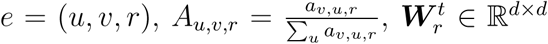 and 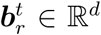 stand for the learnable parameters, *ReLU* (*x*) = *max*(0*, x*), and *N_r_*(*v*) = *{u, u ∈ V, u ≠ v,* (*u, v, r*) *∈ E}* denotes the set of adjacent nodes connected to *v ∈ V* through edges of type *r ∈ R*.

Then the feature 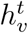 of node *v* is updated by

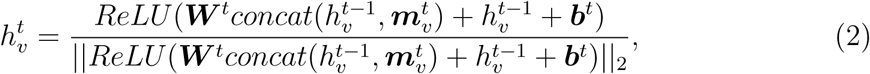

where ***W*** *^t^ ∈* ℝ*^d×d^* and ***b****^t^ ∈* ℝ*^d^* stand for the learnable parameters, and *concat*(*·, ·*) stands for the concatenation operation.

Finally, the confidence score *s_u,v_* of the relation *r* between node *u* and node *v* is derived from the learned node embeddings and the corresponding projection matrices, that is,

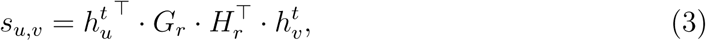

where *G_r_, H_r_ ∈* ℝ*^d×k^* stand for the edge-type specific projection matrices.

We minimized the Bayesian personalized ranking (BPR) loss [71] for drug-target interaction reconstruction, by regarding those edges not in the edge set *E* as missing values rather than negative samples, that is,

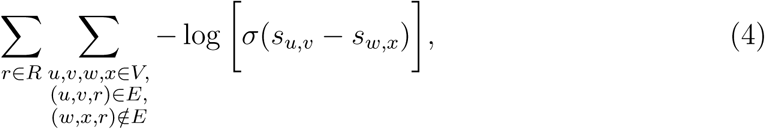

where, *s_u,v_* and *s_w,x_* stand for the confidence scores of the relation *r* between *u* and *v* and between *w* and *x*, respectively, and *σ*(*·*) stands for the sigmoid activation function. Intuitively, in the above loss function, the confidence scores of the node pairs (*u, v*) in the edge set (i.e., (*u, v, r*) ∈ *E*) were encouraged to be higher than those of unseen pairs (*w, x*) (i.e., (*w, x, r*) ∉ *R*).

We predicted the confidence scores under the relation of virus target-drug interactions for each virus target-drug pair using Equation (3). Then the confidence scores were averaged across all the proteins of a certain virus (e.g., SARS-CoV, MERS-CoV or SARS-CoV-2), and the corresponding p-values were obtained by z-test. For each virus, we selected those predictions with a p-value *<* 0.05 as drug candidates.

### 4.3. Automated relation extraction from large-scale literature texts

We used a deep learning based relation extraction method named BERE [4] to extract the coronavirus related drugs from large-scale literature texts. More specifically, the sentences mentioning the two entities of interest, i.e., name (or alias) of a coronavirus or coronavirus target, or name (or alias) of a drug, were first collected using a dictionary-based name entity recognition method (string matching). For each pair of entities (*e*_1_*, e*_2_), there are usually more than one sentence describing the underlying relations. Therefore, we used a bag of sentences *S_e_*_1_ *_,e_*_2_, denoting the set of all the sentences mentioning both *e*_1_ and *e*_2_, to predict the relation between these two entities.

We first encoded each sentence *s ∈ S_e_*_1_ *_,e_*_2_ in a semantic and syntactic manner using a hybrid deep neural network (*h*: *s →* ℝ*^d^*), including a self-attention module [72], a bidirectional gated recurrent unit (GRU) module [73] and a Gumbel tree-GRU module [4, 74]. Each sentence representation *h*(*s*) was then scored by a sentence-level attention module to indicate its contribution to the relation prediction, that is,

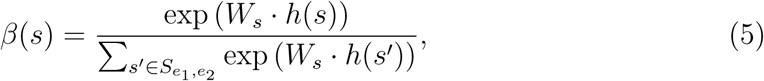

where *β*(*s*) *∈* ℝ stands for the weight score, and *W_s_ ∈* R*^d×^*^1^ stands for the learnable weight parameters. Finally, the relation was predicted by a binary classifier, based on the weighted sum of sentence representations, that is,

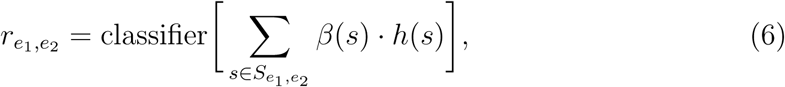

where *r_e_*_1_ *_,e_*_2_ stands for the probability of the relation of interest between entities *e*_1_ and *e*_2_ mentioned by the bag of sentences *S_e_*_1_ *_,e_*_2_.

The training corpus we used was curated automatically from nearly 20 million PubMed (http://www.pubmed.gov) abstracts by a distant supervision technique [75]. In detail, the names (or aliases) of drugs or targets in sentences were first annotated by a dictionary-based named entity recognition method (string matching), in which the name dictionary was derived from DrugBank (version 5.1.0) [17], with ambiguous names (e.g., common words) removed. Next, the label for each bag of sentences co-mentioning a drug-target pair of interest was annotated automatically by the known drug-target interactions in DrugBank. The unlabeled corpus that we used in this work for text-mining the coronavirus related drugs was obtained from approximately 2.2 million PMC full-text articles, with entities of interest annotated using the aforementioned named entity recognition approach. A coronavirus related drug was extracted as a hit candidate if the model found a bag of sentences describing a relation between this drug and a target in the coronavirus of interest.

### 4.4. Connectivity map analysis

We used the transcriptome analysis approach to further filter the potential drug candidates for treating the COVID-19 patients infected by SARS-CoV-2. Due to the lack of gene expression data from the SARS-CoV-2 infected patients, we used those from the SARS-CoV infected patients to screen the potential therapeutic drug candidates against COVID-19. Such a strategy is reasonable as SARS-CoV and SARS-CoV-2 are two closely related and highly similar coronavirus. First, the genome of SARS-CoV-2 is phylogenetically close to that of SARS-CoV, with about 79% of sequence identity [76], and the M (membrane), N (nucleocapsid) and E (envelope) proteins of these two coronaviruses have over 90% sequence similarities [77]. In addition, the pathogenic mechanisms of SARS-CoV-2 and SARS-CoV were highly similar [78].

In particularly, we collected the gene expression profiles of the peripheral blood mononuclear cells (PBMCs) from ten SARS-CoV infected patients (GEO:GSE1739) [6]. The raw gene expression values were first converted into logarithm scale, and then the differential expression values (z-scores) were computed by comparing to those of healthy persons using the same protocol as described in [5], that is,

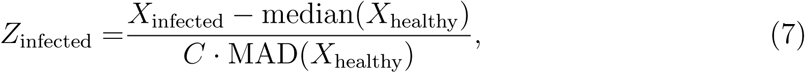

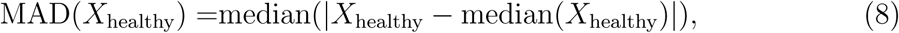

where *Z*_infected_ stands for the z-scores of the SARS-CoV infected patients, *X*_infected_ and *X*_healthy_ stand for the gene expression values in logarithm scale of the infected and healthy persons, respectively, median(*·*) stands for the median operation, MAD(*·*) stands for the median absolute deviation operation, and *C* = 1.4826 is a constant for normalization. The p-values for all the genes with measured expression values during the analysis were also computed based on the z-scores. The up- and down-regulated genes were then identified using a cut-off threshold of p-value *<* 10*^−^*^10^. We used the connectivity map (CMap) [5], which contains the cellular gene expression profiles under the perturbation of 2428 well annotated reference compounds, to measure the associations of gene expression patterns between SARS-CoV infected patients and the reference compound-perturbed cells. The connectivity map scores were computed based on the up- and down-regulated gene sets of SARS-CoV infected patients using the web tool (https://clue.io/query). Under the hypothesis that the gene expression pattern resulting from the perturbation by a therapeutic compound should be negatively correlated with that resulting from the coronavirus infection, we selected those compounds that have significant negative connectivity map scores, that is, the list of drug candidates predicted to treat the coronavirus infected patients was obtained by selecting the compounds with the connectivity map scores *< −*90, which was suggested by the original paper [5].

### 4.5. Cells and virus

The African green monkey kidney Vero E6 cell line was purchased from the Cell Resources Center of Shanghai Institute of Life Science, Chinese Academy of Sciences (Shanghai, China) and cultured in DMEM medium (Gibco Invitrogen, no. 12430-054) containing 10% fetal bovine serum (FBS; Gibco Invitrogen) at 37 *^◦^*C with 5% CO_2_ atmosphere. BetaCoV/JS03/human/2020 (EPI ISL 411953), a SARS-CoV-2 virus strain, was isolated from nasopharyngeal swab of a 40-year old female confirmed as COVID-19 case by reverse transcriptase polymerase chain reaction (RT-PCR) in December 2019.

The virus was propagated in Vero E6 cells, and the viral titer was determined by the 50% tissue culture infective dose (TCID50) based on microscopic observation of cytopathic effects. All the *in vitro* SARS-CoV-2 infection experiments were performed in a biosafety level-3 (BLS-3) laboratory in Jiangsu Provincial Center for Diseases Control and Prevention, Jiangsu, China.

### 4.6. Antiviral drugs

Potential antiviral drugs, including zanamivir, oseltamivir, remdesivir, baricitinib, olaparib and arbidol, were all provided by MCE (Medchem Express, China). The PARP1 inhibitor mefuparib hydrochloride (CVL218) with a purity of more than 99.0% was provided by Convalife, Shanghai, China.

### 4.7. Cytotoxicity test and virus infection assay

The cytotoxicity of the tested drugs on Vero E6 cells was determined by the CCK8 assays (Beyotime, China). At 48 h post addition of the tested drugs, 20 *µ*L CCK8 was added to each well and incubated at 37 *^◦^*C for 1 h. Then optical density was measured at 450 nm. The 50% cytotoxic concentration (CC_50_) values were calculated using GraphPad Prism (GraphPad Software, USA). Vero E6 cells were seeded into 96-well plates with a density of 5 *×* 10^4^ cells/well for incubation in DMEM medium supplemented with 10% FBS for 16 h in an incubator with 5% CO_2_ at 37 *^◦^*C, for cells to reach 80% confluent. Then, cell culture medium of each well was removed, and PBS was used to wash the cells once, before evaluating the antiviral efficacy of the drugs. Four duplicated wells were made for each dose of drugs, and the cells were pre-treated with different doses of antiviral drugs diluted by the cell maintenance solution (50 *µ*L per well) for 1 h. For the virus control and cell control wells, cell medium containing DMSO or only medium of 50 *µ*L per well was added. Next, pre-treated or untreated cells in each well were infected with the virus with multiplicity of infection (MOI) of 0.05 for 2 h. After that, the virus-drug mixture was removed and cells were further cultured with fresh drug-containing medium at 37 *^◦^*C with 5% CO_2_ atmosphere for 48 h. Then culture supernatant per well was harvested and inactivated at 56 *^◦^*C for 30 min to further extract and quantify viral RNA.

### 4.8. Viral RNA extraction and quantitative real-time PCR (qRT-PCR)

Viral RNA was extracted from culture supernatant using the HP RNA Isolation Kit (Roche) according to the manufacturer’s instructions. RNA was eluted in 30 *µ*L RNase-free water. Reverse transcription was performed with a SARS-CoV-2 nucleic acid detection kit (BioGerm, China) according to the manufacturer’s instructions. The PCR reaction system was configured as follows: 6 *µ*L of qRT-PCR reaction solution, 2 *µ*L of qRT-PCR enzyme mixture, 2 *µ*L of primer probe and 2.5 *µ*L of template, and the reaction was performed as follows: 50 *^◦^*C for 10 min, 95 *^◦^*C for 5 min, followed by 40 cycles of 95 *^◦^*C for 10 s, 55 *^◦^*C for 40 s. The values of 2*^−^*^Δ^*^CT^* were calculated according to the CT value measured from the PCR instrument, to represent the relative virus copies of the drug group to the control group. The virus replication inhibition rate (%) was calculated as (1*−*2*^−^*^Δ^*^CT^*)*×*100%. The dose-response curves were plotted according to viral RNA copies and the drug concentrations using GraphPad Prism (GraphPad Software, USA).

### 4.9. Time-of-addition assay

To facilitate the observation of the antiviral effects of drugs against SARS-CoV-2 at different timing, relative high doses of the tested drugs (CVL218 at 20 *µ*M and remdesivir at 10*µ*M) were used for the time-of-addition assay. Vero E6 cells with a density of 5 *×* 10^4^ cells per well were treated with the tested drugs, or DMSO as controls at different stages of virus infection. The cells were infected with virus at an MOI of 0.05. The “Full-time” treatment was to evaluate the maximum antiviral effects, with the tested drugs in the cell culture medium during the whole experiment process, which was consistent with the descriptions in the virus infection assay. For the “Entry” treatment, the tested drug was added to the cells for 1 h before virus infection, and then cells were maintained in the drug-virus mixture for 2 h during the virus infection process. After that, the culture medium containing both virus and the tested drug was replaced with fresh culture medium till the end of the experiment. For the “Postentry” experiment, virus was first added to the cells to allow infection for 2 h before the virus-containing supernatant was replaced with drug-containing medium until the end of the experiment. At 14 h post infection, the viral inhibition in the cell supernatants of the tested drug was quantified by qRT-PCR, and calculated using the DMSO group as reference.

### 4.10. Indirect immunofluorescence assay

Vero E6 cells were treated with CVL218 at 5 *µ*M, 15 *µ*M and 25 *µ*M, respectively, following the same procedure of “full-time” treatment. Infected cells were fixed with 80% acetone in PBS and permeabilized with 0.5% Triton X-100, and then blocked with 5% BSA in PBS buffer containing 0.05% Tween 20 at room temperature for 30 min. The cells were further incubated with a rabbit polyclonal antibody against a SARS-CoV nucleocapsid protein (Cambridgebio, USA) as primary antibody at a dilution of 1:200 for 2 h, followed by incubation with the secondary Alexa 488-labeled goat anti-rabbit antibody (Beyotime, China) at a dilution of 1:500. Nuclei were stained with DAPI (Beyotime, China). Immunofluorescence was observed using fluorescence microscopy.

### 4.11. Western blot assay

NP expression in infected cells was analyzed by Western blot. Protein samples were separated by SDS-PAGE and then transferred onto polyvinylidene difluoride membranes (Millipore, USA), before being blocked with 6% Rapid Block Buff II (Sangon Biotech, China) at room temperature for 10 min. The blot was probed with the antibody against the viral nucleocapsid protein (Cambridgebio, USA) and the horseradish peroxidase-conjugated Goat Anti-Rabbit IgG (Abcam, USA) as the primary and the secondary antibodies, respectively. Protein bands were detected by chemiluminescence using an ECL kit (Sangon Biotech, China).

### 4.12. CpG-PDN1826 induced IL-6 production in PBMCs

Peripheral blood mononuclear cells (Yicon, China) were cultured at 37 *^◦^*C at concentration 5% CO_2_ atmospheric on a 96-well plate in RPMI1640 cell growth medium (Corning, Cat.10-040-CVR). For stimulation, PBMC cells were incubated with 1 *µ*M CpG-ODN1826 (InvivoGen, Cat. tlrl-1826). To test whether CVL218 can inhibit IL-6 production, 1 *µ*M and 3 *µ*M concentrations of CVL218 were added to cell culture medium for 6 and 12 h, respectively. The concentration of IL-6 was determined by ELISA using a commercial kit (Dakewe Biotech, Cat. 1110602).

### 4.13. Pharmacokinetics and toxicity study

Sprague-Dawley rats were purchased from Shanghai Laboratory Animal Center, China. The animals were grouped and housed in wire cages with no more than six per cage, under good laboratory conditions (temperature 25 *±* 2*^◦^*C; relative humidity 50 *±* 20%) and with dark and light cycle (12 h/12 h). Only healthy animals were used for experimental purpose. The pharmacokinetics and biodistribution study in Sprague-Dawley rats was approved by Center for Drug Safety Evaluation and Research, Shanghai Institute of Materia Medica, Chinese Academy of Sciences. A total of 144 Sprague-Dawley rats with each sex were used for toxicity study. Animals were randomly separated into four groups (18/sex/group). CVL218 was administered at doses of 20, 40, 60 and 160 mg/kg. For all the groups, 20 rats (10/sex/group) were randomly selected and euthanized at day 28, and their sections of various tissues and organs were obtained and frozen. Ten (5/sex/group) animals were euthanized after a 28-day drug free period, and their sections of tissues and organs were obtained and frozen. Six (3/sex/group) were euthanized after the blood-samples were obtained. For pharmacokinetic and toxicity evaluation, clinical symptoms, mortality and the animals’ body weight were examined. Serum (0.5 mL) was collected to analyze toxicokinetics at different time points post drug administration. The plasma concentration-time data were analyzed using a non-compartmental method (Phoenix, version 1.3, USA) to derive the pharmacokinetic parameters.

### 4.14. Biodistribution study

Thirty Sprague-Dawley rats were randomly divided into three time point groups (3/sex/group). At 3, 6 and 8h after CVL218 administration, animals were sacrificed, and the brain, heart, lung, liver, spleen, stomach and kidney tissues were collected. Tissue samples were washed in ice-cold saline, blotted with paper towel to remove excess fluid, and weighed. Tissue samples were fluid, weighted and stored at *−*20 *±* 2 *^◦^*C until the determination of drug concentration by LC-MS-MS.

### 4.15. Toxicity study in cynomolgus monkeys

Healthy male and female cynomolgus monkeys aged 3–4 years were purchased from Guangdong Landau Biotechnology, China. The animals were maintained in accordance with the Guide for the Care and Use of Laboratory Animals.

Cynomolgus monkey (5/sex/group) were selected using a computerized randomization procedure, and administered CVL218 by nasogastric feeding at dose levels of 0 (control), 5, 20, 80 mg/kg. Individual dose volumes were adjusted weekly based on body weight of monkeys. The monkeys were observed twice daily for viability/mortality and for any change in behavior, reaction to treatment or ill-health. Electrocardiograms, intraocular pressure, rectal temperature and body weight were recorded. For all the groups, 2/3 of the animals were randomly selected and euthanized at day 28. The remaining animals were euthanized after a 28-day drug free period. Blood samples were taken before and at 0.5, 1, 2, 4, 8 and 24 h post-dose on days 1 and 28 of the treatment period. Pharmacokinetic evaluation was performed using a non-compartmental method (Phoenix, version 1.3, USA) and pharmacokinetic parameters were calculated for individual monkeys.

### 4.16. Statistical analysis

All data represent the means *±* standard deviations (SDs) of n values, where n corresponds to the number of data points used. The figures were prepared using GraphPad Prism (GraphPad Software, USA). The statistical significance was calculated by SPSS (ver.12), and two values were considered significantly different if the p-value is *<* 0.05.

### 4.17. Molecular docking

The docking program AutoDock4.2 [79] was used to model the molecular interactions between PARP1 inhibitors CVL218 and olaparib to the N-terminal domain of the N protein of SARS-CoV-2 (SARS-CoV-2-N-NTD). The structure of SARS-CoV-2-N-NTD used for molecular docking was built from homology modeling [80]. The AutoGrid program was used to generate a grid map with 60*×*60*×*60 points spaced equally at 0.375 Å for evaluating the binding energies between the protein and the ligands.

## Acknowledgement

This work was supported in part by the National Natural Science Foundation of China (61872216, 81630103, 31900862), Jiangsu Provincial Emergency Project on Prevention and Control of COVID-19 Epidemic (BE2020601), the Nation Science and Technology Major Projects for Major New Drugs Innovation and Development (2018ZX09711003-004-0022019ZX09301010), Pudong New Area Science and Technology Development Foundation (PKX2019-S08), and the Turing AI Institute of Nanjing, and the Zhongguancun Haihua Institute for Frontier Information Technology. The authors thank Dr. Feixiong Cheng for email communications on the connectivity map analysis.

## Supplementary Materials

### Detailed docking results on the interactions between PARP1 inhibitors and the N-terminal domain of the nucleocapsid protein of SARS-CoV-2

We noticed that the binding surface of HCoV-OC43-N-NTD with PJ-34 consists of the N-terminal loop, *β*2 and *β*3 strands (Figure 5B left panel). The oxygen and nitrogen atoms on the 6-phenanthridinone of PJ-34 form three hydrogen bonds with S64 (3.1 Å), Y126 (3.0 Å) and F66 (water-mediated) of HCoV-OC43-N-NTD. In addition to the hydrogen network, the aromatic ring of phenanthridinone on PJ-34 participates in *π*-stacking with H104 on *β*2 strand and Y124 on *β*3 strand. Compared to CoV-OC43-N-NTD, the binding pocket of SARS-CoV-2-N-NTD encompassed by *β*2 strand, *β*3 strand and loops is approximately similar in structural compositions, but more spacious, which may facilitate to bind with larger molecules. As shown in the Figure 5B (middle panel), CVL218 can be reliably docked inside the pocket of SARS-CoV-2-N-NTD with key residues including Y111 on *β*2 strand, R92 on *β*3 strand, as well as S51 and E118 on loops. Among them, Y111 forms a bifurcated hydrogen bond to the oxygen atom of the benzofuran ring and the nitrogen atom of the amide group on CVL218 with a distance of 2.6 Å and 3.1 Å, respectively. In addition, R92 further stabilizes the CVL218 molecule by forming a hydrogen bond with the oxygen atom of the amide group on the benzofuran ring. The nitrogen atom of the amine close to the benzene ring of CVL218 also forms a hydrogen bond of distance 2.8 Å with the side chain of E118. In addition to the hydrogen network which plays an essential role in CVL218 binding, the hydrophobic interactions involved by T49, Y109, and Y112 of SARS-CoV-2-N-NTD also contribute to the molecular interaction.

The binding surface of olaparib on SARS-CoV-2-N-NTD is similar but not exactly the same to that of CVL218. The two carbonyl groups of olaparib form hydrogen bonds with residues S51 and R149 of SARS-CoV-2-N-NTD with distances 3.0 Å and Å, respectively. In addition, the residues around the binding surface (i.e., R88, R92, Y109, Y111, R149, P151) participate in the interaction with olaparib via hydrophobic interactions.

**Figure S1:**
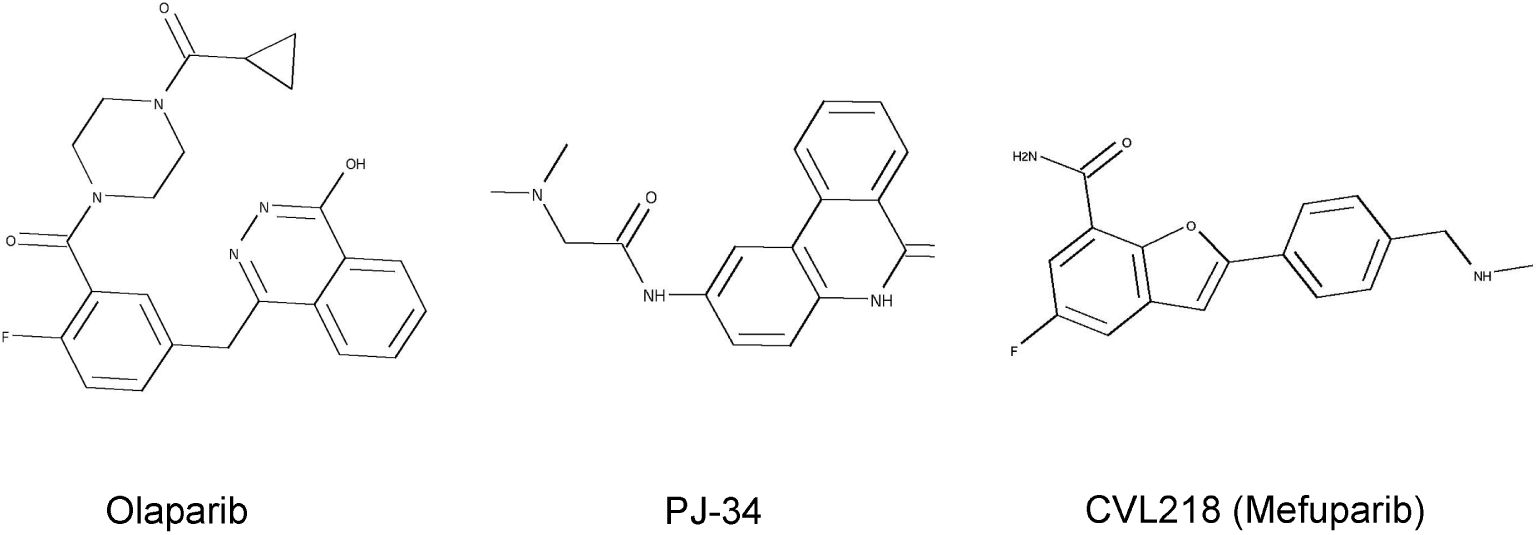
Structures of PARP1 inhibitors mentioned in this study.

**Figure S2:**
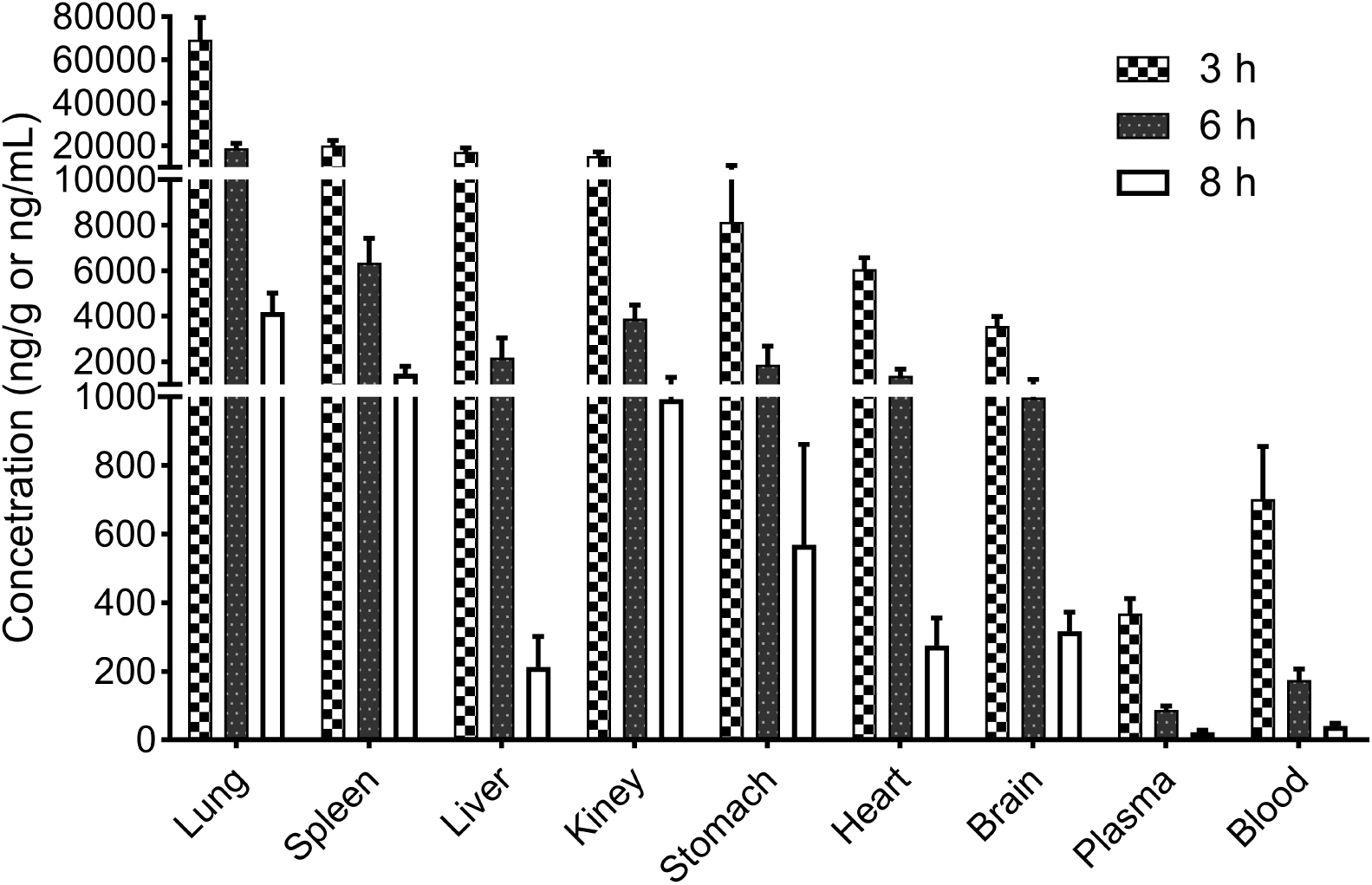
Tissue distribution characteristics of CVL218 in rats, with the highest concentration in lung. The concentrations of CVL218 in different tissues were measured at the 3/6/8 h time points after 20 mg/kg oral administration to rats. With the extension of administration time, the concentration of CVL218 in each organ decreased in a time-dependent manner.

**Table S1:** The top list of perturbational classes identified by the connectivity map analysis using the gene expression profiles of the peripheral blood mononuclear cell (PBMC) samples of ten SARS-CoV-infected patients [6].

| Connectivity map score | Perturbational classes |
| --- | --- |
| −40.84 | **PARP inhibitor** |
| −37.31 | RNA Polymerase Enzymes LOF |
| −37.27 | DNA synthesis inhibitor |
| −36.55 | GABA receptor antagonist |
| −29.62 | MDM inhibitor |

**Table S2:** Comparison of the tissue distributions of CVL218 and arbidol in rats, following 20 mg/kg and 54 mg/kg oral administrations, respectively.

| Drugs | Dose<br>(mg/kg) | Time<br>(min) | Lung | Spleen | Liver | Kidney | Stomach | Heart | Brain |
| --- | --- | --- | --- | --- | --- | --- | --- | --- | --- |
| CVL218 | 20 | 180 | 69318±10476 | 20202±2300 | 17215±1919 | 15201±1984 | 8145±2624 | 6077±496 | 3580±416 |
|  |  | 360 | 18858±2365 | 6358±1058 | 2187±859 | 3903±594 | 1871±813 | 1390±292 | 998±220 |
|  |  | 480 | 4183±847 | 1475±324 | 213±88 | 993±327 | 569±293 | 275±80 | 317±55 |
| Arbidol<sup>a</sup> | 54 | 5 | 933±837 | 48±35 | 104±82 | 79±54 | 8210±5410 | 72±47 | 101±67 |
|  |  | 15 | 2603±1848 | 519±281 | 963±290 | 259±190 | 23180±10170 | 132±69 | 50±10 |
|  |  | 360 | 833±397 | 143±51 | 262±175 | 58±21 | 52750±3059 | 41±28 | 31±21 |

a. The concentrations of arbidol in different tissues of rats at 5/15/360 min time points with 54 mg/kg oral administration were obtained from [92].

**Table S3:** Comparisons of pharmacokinetic parameters in rats between CVL218 and arbidol following 20/40 mg/kg and 18/54 mg/kg oral administrations, respectively.

| Drugs | Dose<br>(mg/kg) | Gender | Tmax<br>(h) | Cmax<br>(ng/mL) | AUC<sub>0-t</sub><br>(ng·h/mL) | AUC<sub>0-∞</sub><br>(ng·h/mL) | MRT<sub>0-∞</sub><br>(h) | t<sub>1/2</sub><br>(h) |
| --- | --- | --- | --- | --- | --- | --- | --- | --- |
| CVL218 | 20 | male | 4.0 (4.0$\tilde{4}$.0) | 234 ± 35 | 1070 ± 176 | 1111 ± 192 | 3.91 ± 0.19 | 1.19 ± 0.09 |
| | | female | 3.0 (2.0$\tilde{3}$.0) | 502 ± 80 | 2196 ± 228 | 2222 ± 241 | 3.16 ± 0.41 | 1.1 ± 0.16 |
| | | total | 3.5 (2.0$\tilde{4}$.0) | 368 ± 157 | 1633 ± 643 | 1666 ± 639 | 3.54 ± 0.50 | 1.15 ± 0.13 |
| | 40 | male | 3.0 (2.0$\tilde{4}$.0) | 510 ± 259 | 2802 ± 967 | 2830 ± 983 | 4.51 ± 0.18 | 1.3 ± 0.33 |
| | | female | 2.0 (2.0$\tilde{3}$.0) | 940 ± 117 | 5220 ± 1113 | 5242 ± 1115 | 4.05 ± 0.43 | 1.29 ± 0.21 |
| | | total | 2.5 (2.0$\tilde{4}$.0) | 725 ± 296 | 4011 ± 1620 | 4036 ± 1620 | 4.28 ± 0.39 | 1.3 ± 0.24 |
| Arbidol<sup>a</sup> | 18 | male | 0.28 ± 0.11 | 1002 ± 298 | 1956 ± 895 | 2224 ± 1058 | - | 3.6 ± 1.2 |
| | 54 | male | 0.18 ± 0.06 | 4711 ± 2361 | 6790 ± 2749 | 7558 ± 2877 | - | 3.3 ± 0.7 |

a. The pharmacokinetic data of arbidol were obtained from [91].

**Table S4:**
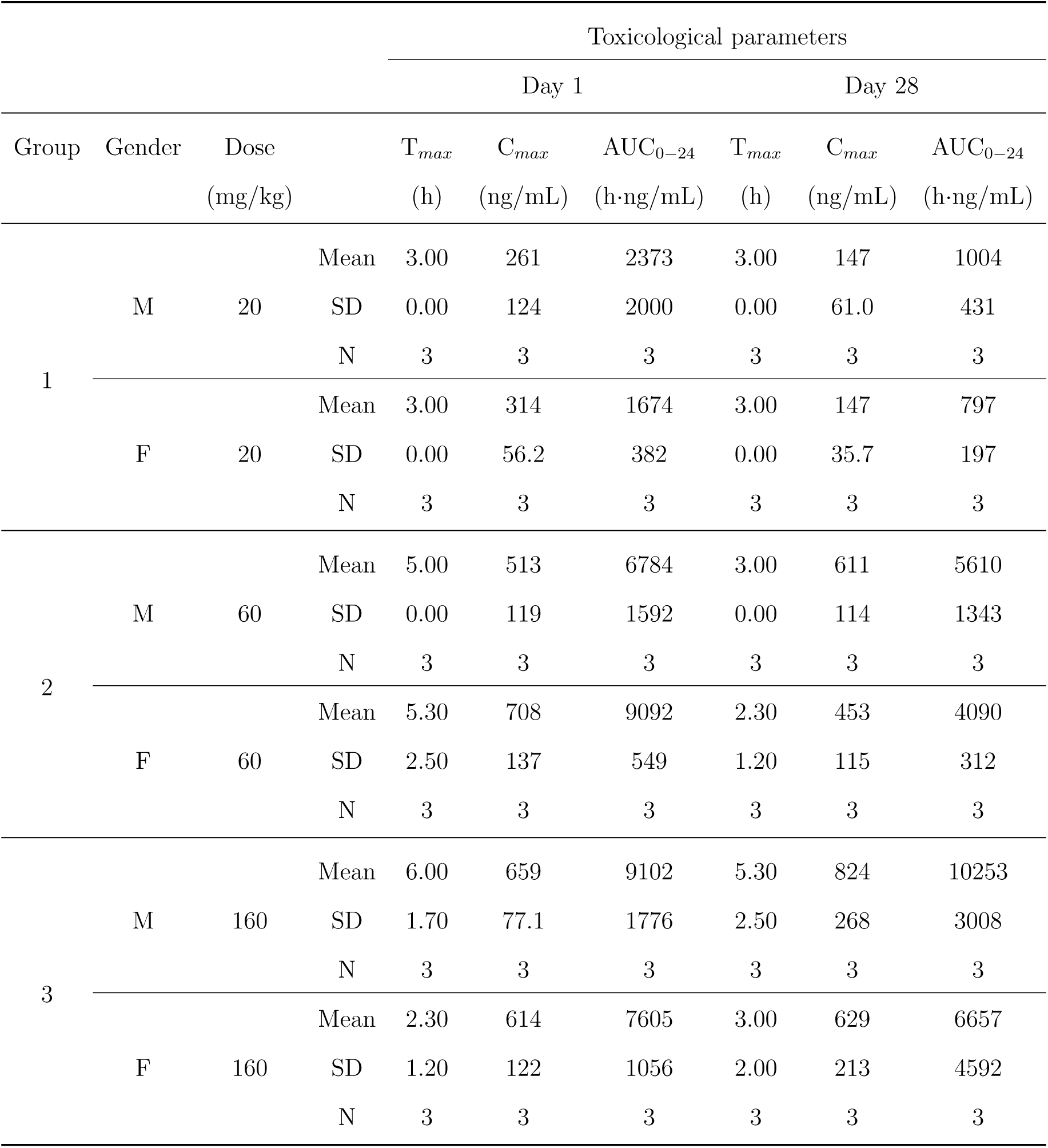
Toxicokinetic parameters of CVL218 in rats in a four-week toxicity study.

**Table S5:**
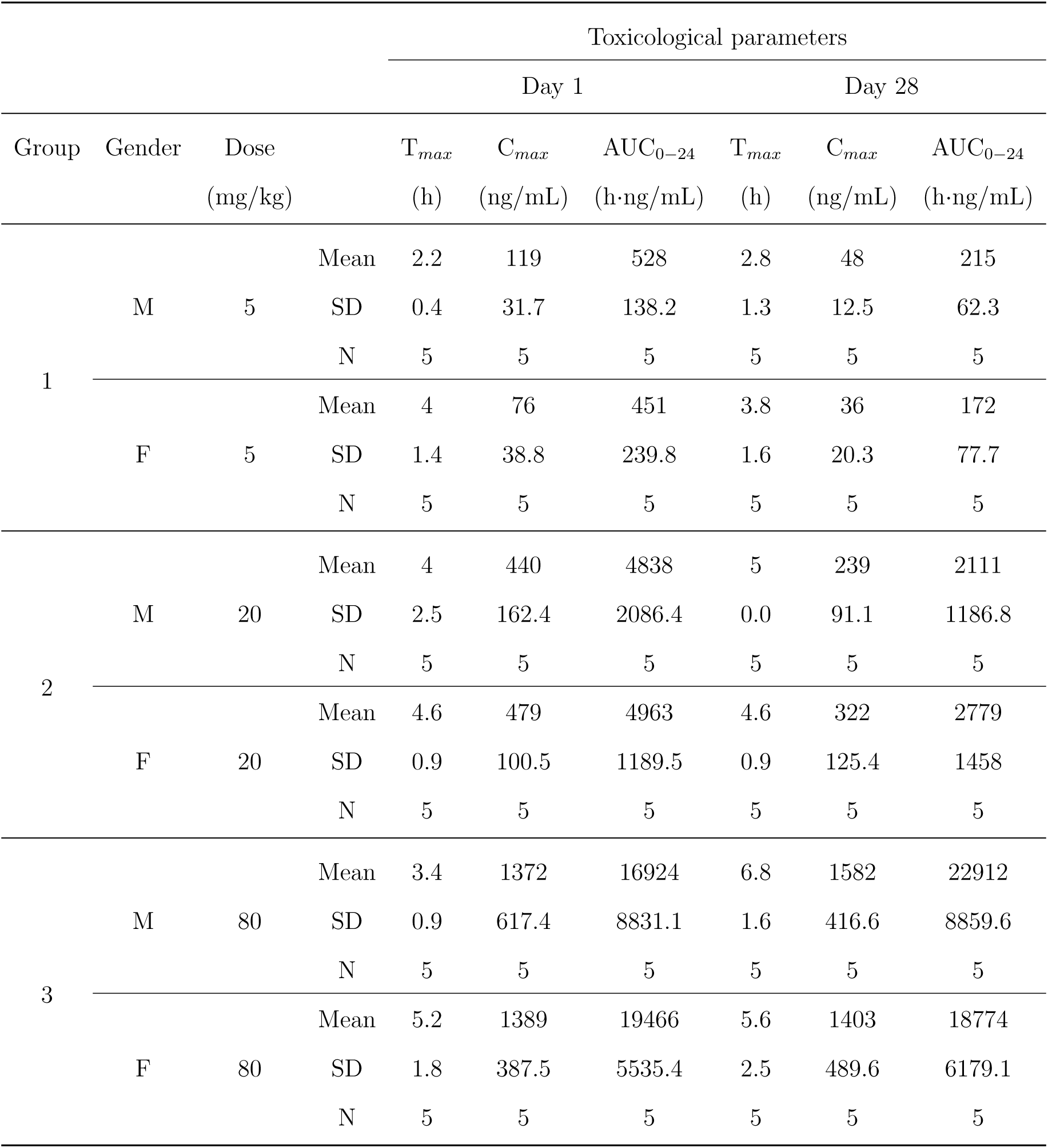
Toxicokinetic parameters of CVL218 in monkeys in a four-week toxicity study.

